# Circadian regulators PER1 and PER2 regulate osteoclastogenesis by balancing competing activities of innate immunity genes

**DOI:** 10.64898/2026.01.11.698889

**Authors:** Nobuko Katoku-Kikyo, Elizabeth K. Vu, Samuel Mitchell, Ismael Y. Karkache, Elizabeth W. Bradley, Nobuaki Kikyo

**Author notes:** Corresponding authors: (EWB) and (NK).

## Abstract

Bone remodeling is influenced by circadian rhythms as demonstrated by global gene expression patterns and the phenotypes of knockout mice of circadian regulators. However, the direct connections between circadian regulators and specific bone genes remain unclear. We previously found that a conditional knockout of *Per1*, a central circadian regulator, in osteoclasts increased osteoclastogenesis and decreased bone mass, whereas *Per2* knockout did not cause these phenotypes. Here, we extended the research to *Per1*;*Per2* conditional double knockout mice and observed different phenotypes and underlying mechanisms from individual knockouts. In contrast to *Per1* knockout, the double knockout decreased osteoclastogenesis and increased bone mass. This was accompanied by downregulation of genes involved in innate immunity, including several known promoters and inhibitors of osteoclastogenesis. Chromatin immunoprecipitation and reporter assay suggested direct regulation of some of them by PER proteins. These results indicate that PER1 and PER2 are critical regulators of osteoclastogenesis through balancing multiple competing activities of osteoclastogenesis, rather than acting as simple promoters or inhibitors of osteoclastogenesis. Regulation of innate immunity genes by circadian regulators is widely observed across other monocyte/macrophage lineages. Our results extend this common mechanism to osteoclasts with therapeutic potential to treat inflammatory bone diseases.

**Lay summary:** Although circadian rhythms regulate bone remodeling, direct links between circadian regulators and bone genes remain unclear. We previously demonstrated that depletion of *Per1*, a main circadian regulator, downregulates immunological genes, increases the number of osteoclasts, the main bone resorbing cells, and decreases bone mass in mice. Here, we showed the opposite effects with double depletion of *Per1* and *Per2* and identified a new set of downregulated immunological genes that promote or inhibit osteoclastogenesis. This study connects circadian rhythms to bone resorption through immunological genes with opposing activities in osteoclastogenesis, supporting therapeutic interventions targeting circadian regulators to treat inflammatory bone diseases.

## Introduction

Circadian rhythms of the body align many aspect of physiology to the environmental light-dark cycle with a period of 24 h. Bone remodeling is no exception.^1,2^ More than 25% of the transcriptome in the mouse bone is under circadian control, including those encoding the master regulators for osteoblastogenesis (*Runx2*) and osteoclastogenesis (*Ctsk*), as well as structural (e.g., procollagens) and signaling proteins (e.g., bone morphogenic proteins and colony stimulating factor 1). Although such data is not available in humans, serum and urine concentrations of bone turnover markers (various collagen fragments) have been used as alternative markers for circadian rhythmicity. In addition, other parameters related to bone metabolism—serum concentration of parathyroid hormone, calcitonin, calcium ion, and phosphate—display circadian fluctuations. The functional role of circadian rhythms in bone remodeling is exemplified by the increases the risk of osteoporosis in shift workers. However, the relationship between specific circadian regulators and bone genes remains unclear.^3,4^.

Circadian rhythms in the bone are, just like in virtually any other tissues, driven by the BMAL1-CLOCK heterodimer as the central regulator.^5,6^ This complex generally activates hundreds of target genes by binding to the E box (CANNTG) in their promoters and enhancers. The target genes include the repressors *Cry* (*Cry1* and *Cry2*) and *Per* (*Per1*, *Per2*, and *Per3*). The repressors inhibit the BMAL1-CLOCK complex through direct binding, downregulating the target genes; to date, *Per3*’s roles remain unclear. Several hours later, BMAL1-CLOCK is released from this inhibition due to the degradation of CRY and PER via phosphorylation and ubiquitination. This feedback loop is completed with a period of 24 h. Additional feedback loops imposed by the BMAL1-CLOCK target *Ror* (retinoic acid receptor-related orphan receptor) and *Nr1d* (Rev-erbs or reverse orientation c-erbA), which are not directly involved in the current study.

Phenotypic analysis of bone cell-specific conditional knockout (cKO) mouse models should have provided direct evidence of the circadian control of bone remodeling.^1,2^ However, published results are largely limited to *Bmal1* KO and their phenotypes are not clear. To address this limitation, we previously developed *Per1* cKO mice and *Per2* cKO mice using *Cx3cr1* promoter-driven *Cre*, which is mainly active in the monocyte/macrophage lineages, including osteoclasts.^7^ *Per1* cKO increased osteoclastogenesis and diminished bone mass, accompanied by downregulation of seven inflammatory genes known to increase or decrease osteoclastogenesis upon depletion. A current interpretation is that the cumulative effects of their activities promote osteoclastogenesis, leading to the decreased bone mass.

To understand whether *Per1* and *Per2* have redundant roles in osteoclastogenesis not revealed by single KO, we developed *Per1*;*Per2* double cKO mice (dKO) in the current study. Unexpectedly, dKO mice showed the opposite phenotypes to *Per1* cKO mice. Further, RNA-seq, chromatin immunoprecipitation (ChIP), and reporter assays indicated that PER1 and PER2 redundantly regulated genes involved in innate immunity that acted as promoters or inhibitors of osteoclastogenesis. These findings support the idea that PERs regulate osteoclastogenesis by balancing opposing activities involved in immune responses.

## Materials and Methods

### Generation of double cKO mice of *Per1* and *Per2*

*Cx3cr1* promoter-driven *Per1* cKO mice (*Cx3cr1-Cre^tg/wt^;Per1^fl/fl^*) and *Per2* cKO mice (*Cx3cr1-Cre^tg/wt^;Per2^fl/fl^*) were described before.^7^ These mice were mated to create dKO mice (*Cx3cr1-Cre^tg/wt^;Per1^fl/fl^;Per2^fl/fl^*) and littermate control called Cont (*Cx3cr1-Cre^wt/wt^;Per1^fl/fl^;Per2^fl/fl^*). Genotypes of the mice were verified as described before. The mice were housed under a 12 h light-12 h dark cycle with water and food provided *ad libitum*. All mouse procedures were approved by the Institutional Animal Care and Usage Committee of the University of Minnesota (2410-42491A). All mouse experiments comply with the Guide for the Care and Use of Laboratory Animals published by the US National Institute of Health.

### Micro-CT scan analysis of the bones

Femurs were applied to micro-CT scanning as previously described.^7^ More specifically, femurs of 12-week-old mice were fixed in 10% formalin in phosphate-buffered saline (PBS) for 24 h and stored in 70% ethanol at 4°C. Micro-CT images were captured with an XT H 225 CT scanning system (Nikon Metrology) with the following settings: 120 kV, 61 μA, 17 min/scan, 720 projections at two frames/projection with an integration time of 708 ms, and an isometric voxel size of 7.11 μm using a 1 mm aluminum filter. Bone morphometric analysis was performed with SkyScan CT Analyzer (CTAn, Brucker micro-CT) and the results were indicated with standard symbols.^8^ The 3D bone images were reconstructed with CT Pro 3D program (Nikon Metrology), converted to bitmap data with VGStudio MAX 3.2 (Volume Graphics), and reoriented with DataViewer (SkyScan, Buker micro-CT) for qualitative analysis. 3D analysis of trabecular bones was performed at the distal metaphysis 0.7 mm proximal to the growth plate, extending 1.5 mm toward the diaphysis. 3D analysis of cortical bones was performed in a 0.5 mm section in the mid-diaphysis 4 mm proximal to the growth plate. Ten femurs were applied to micro-CT in each group and all of them were used in the analysis without exclusion. This number was determined with a preliminary assessment with a power analysis on G*Power (https://download.cnet.com/g-power/3000-2054_4-10647044.html#google_vignette) with a significance level of 0.05 and a power value of 0.8. Bone samples were blinded but not randomized.

### Histomorphometry of bone sections

Tibiae prepared as described above were decalcified in 15% EDTA for 14 days, embedded in paraffin, and sectioned with a thickness of 7 μm. The TRAP staining kit (Sigma Aldrich, 387A-1KT) and Masson’s trichrome staining kit (Polyscience, 25088) were used to stain osteoclasts and osteoblasts, respectively. The numbers of these cells were counted and the ratios between the cells and bone perimeter were calculated using ImageJ. Randomly selected seven tibiae were stained and all of them were applied to the analysis without blinding. This number was determined with G*Power as described above.

### Osteoclast differentiation in vitro

Osteoclasts were prepared from bone marrow monocytes and macrophages (BMMs) as previously described.^7^ On day 1, bone marrow cells were flushed from femurs and tibiae of 8- to 12-week-old mice with PBS and centrifuged at 190 x *g* for 5 min. Cells were resuspended with 1X Red Blood Cell Lysis Buffer (eBioscience, 00-4333-57) and incubated at 25°C for 5 min. Cells were centrifuged again under the same condition and resuspended with Monocyte Medium (25 ng/ml M-CSF [Shenandoah, 100-03], 10% heat-treated fetal bovine serum [FBS], and phenol red-free αMEM [Thermo Fisher, 41061-029]). Cells were incubated with 5% CO_2_ at 37°C overnight. On day 2, non-adherent cells were harvested and applied to a 70 μm cell strainer (Falcon, 352350) to remove cells clusters. They were centrifuged, resuspended in Osteoclast Medium (Monocyte Medium with 100 ng/ml RANKL [Cell Signaling Technology, 68495]), and seeded at 8x10^5^ cells/400 μl/well in a 48-well plate. On day 4, 200 μl of fresh medium was added. On day 5, the medium was replaced with 400 μl of medium. On day 6, the cells were fixed with 4% paraformaldehyde in PBS for TRAP staining. The total number of osteoclasts (TRAP-positive cells with more than two nuclei) in each well were manually counted. ImageJ was used to measure the size of osteoclasts.

### Synchronization of circadian rhythms

Circadian rhythms of osteoclasts were synchronized as previously reported.^7^ On day 5 of differentiation, cells were treated with Synchronization Medium (50% heat-inactivated horse serum in Osteoclast Medium instead of FBS) for 1 h with 5% CO_2_ at 37°C, washed with PBS at 37°C twice, and cultured with fresh Osteoclast Medium.^9^ The time of the completion of these procedures was defined as 0 h post-synchronization. The cells were cultured at 37°C with 5% CO_2_ and harvested for qRT-PCR every 4 h for 48 h (two circadian cycles) starting at 24 h post-synchronization to wait for the recovery from the serum shock following a guideline.^10^ Circadian rhythmicity, amplitude, and period of the PCR values were examined with harmonic regression using Cosinor.Online (https://cosinor.online/app/cosinor.php).^11^

### Bone resorption assay

Bone resorption assay was done following an established protocol.^7^ On day 2 of the osteoclast differentiation, cells were seeded on a bone slice (Immunodiagnostic Systems, DT-1BON1000-96) at 3.2x10^5^ cells/well in a 48-well plate in Osteoclast Medium to induce osteoclastogenesis as described above. The medium was replaced every 3 days after day 6. On day 14, cells were lysed with 10% bleach and the bone slices were washed three times with deionized water. The bone slices were incubated in 200 μl deionized water for 1 h at 25°C twice, stained with Toluidine blue for 30 sec, washed with deionized water three times, and dried. ImageJ was used to quantify stained areas.

### Quantitative RT-PCR (qRT-PCR)

qRT-PCR was performed as described previously.^7^ RNA was extracted from cells with a Quick RNA Microprep kit (Zymo Research, R1051) and applied for cDNA synthesis with ProtoScript II Reverse Transcriptase (New England Biolabs, M0368L). qPCR was done with the primers listed in **Supplementary Table 1** and qPCRBIO SyGreen Blue Mix Lo-ROX (Genesee Scientific, 17-505B). Following PCR conditions were used: initial denaturation at 95°C for 2 min, 40 cycles of 95°C for 5 sec - 60°C for 30 sec - 72°C for 30 sec, and a melting curve step. The levels of mRNA expression were analyzed by normalizing expression values to glyceraldehyde 3-phosphate dehydrogenase (*Gapdh*) expression. Mean ± SEM of biological triplicates with technical triplicates each was calculated.

### Western blotting

Western blotting was done following a published protocol.^12^ Specifically, whole-cell extracts were prepared from 2x10^5^ cells harvested at 36 h and 48 h post-synchronization with an NE-PER Nuclear and Cytoplasmic Extraction kit (Thermo Fisher Scientific, 78833). Proteins were resolved with a 12% SDS-PAGE gel and transferred to an Immobilon P membrane (EMD Millipore, IPVH00010) at 25°C overnight. The next day, the membrane was blocked with 5% non-fat dry milk (BioRad, 180171A) in PBT (0.2% Tween 20 in PBS) for 1 h at 25°C. Proteins were then labeled with the primary antibody diluted in 5% milk in PBT at 25°C for 1 h. After washing with PBT for 5 min three times, the membranes were incubated with secondary antibody in 5% milk in PBT for 1 h at 25°C. After washing with PBT six times, the chemiluminescence signal was detected with a SuperSignal West Dura kit (Thermo Fisher Scientific, 34075) and an iBright Imaging System (Thermo Fisher Scientific). Following antibodies were used: BMAL1 (1/400 dilution, Abcam, ab3350), histone H2B (1/1000, Thermo Fisher Scientific, MA5-14835), goat anti-mouse IgG-HRP (1/200, Bio-Rad, 170-6516), and goat anti-rabbit IgG-HRP (1/200, Bio-Rad, 170-6515).

### Identification of BMAL1-binding sites with published data sets

Following chromatin immunoprecipitation (ChIP)-seq data were downloaded from the Gene Expression Omnibus (GEO) website (https://www.ncbi.nlm.nih.gov/geo/) for **Figure 5A**. A-Input (GSM5044439, mouse liver at ZT4), A-BMAL1 (GSM5044447), B-Input (GSM5044444, mouse liver at ZT4), B-BMAL1 (GSM5044464), C-Input (GSM3885076, mouse liver at ZT8), and C-BMAL1 (GSM3885067). They were uploaded into Integrative Genomics Viewer (IGV, https://igv.org/) via ChIP-Atlas (https://chip-atlas.org/peak_browser) to visualize BMAL1 peaks. Light was on between ZT0 (zeitgeber time 0)–ZT12 and off between ZT12–ZT24.

### ChIP-qPCR

ChIP-qPCR was done as reported before.^13^ Briefly, two million cells were treated with 1% paraformaldehyde for 10 min and then with 125 mM glycine. Chromatin was prepared by the sequential treatment with cell lysis buffer (50 mM HEPES pH7.8, 85 mM NaCl, 0.5% NP-40, and cOmplete Mini Protease Inhibitor Cocktail [Sigma Aldrich, 11 836 153 001]) and nuclear lysis buffer (50 mM Tris-HCl pH 8.0, 10 mM EDTA, 1% SDS, and cOmplete protease inhibitor cocktail). Chromatin was sheared to 200–600 bp with a Bioruptor 300 (Diagenode) and incubated with 2 μg BMAL1 antibody described above or normal mouse IgG (Santa Cruz Biotechnology, sc-2025) as control, 2 μl Dynabeads Protein G (Thermo Fisher Scientific, 10004D), and 400 μl dilution buffer (20 mM Tris-HCl pH 8.0, 150 mM NaCl, 2 mM EDTA, 1% Triton X-100, and cOmplete protease inhibitor cocktail) for 16 h at 4°C. DNA was released from the beads in elution solution (0.1 M NaHCO_3_, 1% SDS, and 200 μg/ml proteinase K) and incubated for 2 h at 65°C to reverse crosslink. DNA was purified with a ChIP DNA Clean & Concentrator (Zymo Research, D5205) and applied to qPCR with the primers listed in **Supplementary Table 2**. Results of biological triplicates with technical triplicates were presented as ratios in comparison to the results with normal mouse IgG.

### Mutation analysis of the E boxes in the *Saa3* and *Mx2* mRNAs

A 515 bp DNA fragment between nucleotide #53,974,881–53,975,396 in chromosome 7 (NCBI37/mm9) was chemically synthesized as the wild type of *Saa3* (Saa3-WT) (**Figure 6A**). This contains two BMAL1-binding motifs, S-Motif-1 and S-Motif-2, located upstream of the *Saa3* gene. In addition, three mutant fragments listed in **Figure 6A** were synthesized by mutating each CANNTG sequence. Similarly, a 449 bp DNA fragment between nucleotide #97,757,290–#97,757,739 in chromosome 16 was synthesized as the wild type of *Mx2* (Mx2-WT) along with four mutant fragments (**Figure 6B**). These sequences were inserted into the minimal promoter vector pGL4.23 encoding the firefly luciferase (F-Luc) (Promega, E841A). 293FT cells (Thermo Fisher Scientific, R70007) were transfected with 100 ng of one of the vectors and 20 ng *Renilla* luciferase (R-Luc) (pcDNA3.1dsRluc, Addgene, 68053) vector with Lipofectamine 2000 (Thermo Fisher Scientific, 11668027) as described before.^14^ After 24 h the cells were harvested for the luciferase assay with a Dual-Luciferase Reporter Assay system (Promega, E1910). The transfection efficiency was normalized by calculating the ratio between F-Luc and R-Luc activities. Relative F-Luc/R-Luc ratio of each mutant was calculated by defining the ratio with the WT sequence as 1.0.

### Transfection of circadian genes into RAW264.7 cells

RAW264.7 cells (ATCC, TIB-71) were transfected with empty vector or plasmids encoding *Clock*, *Bmal1*, *Per1*, and *Per2* as follows.^7^ On day 1, cells were seeded at 8x10^4^ cells/well in a 48-well plate in 10% FBS in DMEM. On day 2, total 1.6 μg of the plasmid was transfected with 2 μl Lipofectamine LTX and 1.5 μl PLUS Reagent (Invitrogen, 15338030) following the instructions. The cells were incubated with 5% CO_2_ at 37°C and harvested 48 h later for qRT-PCR.

### RNA-seq

RNA-seq and data analysis were performed as previously described.^7^ Total RNA was prepared from osteoclasts on day 6 and mRNA was purified with poly-T oligo nucleotide attached to magnetic beads to synthesize cDNA with random hexamer primers. Non-directional libraries were prepared and completed by end repair, A-tailing, adapter ligation, size selection, amplification, and purification. The libraries were sequenced on an Illumina platform and paired-end reads were generated. Raw reads of the fastq format were processed through in-house perl scripts. Paired-end clean reads were aligned to the reference genome Mus musculus GRCm38 (ftp://ftp.ensembl.org/pub/release-94/fasta/mus_musculus/dna/Mus_musculus.GRCm38.dna.primary_assembly.fa.gz and ftp://ftp.ensembl.org/pub/release-94/gtf/mus_musculus/Mus_musculus.GRCm38.94.gtf.gz) using Hisat2 v2.0.5 (https://daehwankimlab.github.io/hisat2/). featureCounts v1.5.0-p3 (http://subread.sourceforge.net/) was used to count read numbers mapped to each gene. Differential expression analysis was performed with the DESeq2 R package (1.20.0) (https://www.r-project.org/). Differential gene expression was defined as > 1.5-fold change (log2 fold change > 0.58 or < –0.58) and an adjusted p-value < 0.05. Enriched gene pathways in differentially expressed genes were identified with the clusterProfiler R package (https://www.r-project.org/) and Gene Ontology (GO, http://www.geneontology.org). Adjusted p-value < 0.05 was considered as significant enrichment.

### Gene network analysis

The STRING program (https://string-db.org/) was applied to investigate the interactome of the proteins and genes involved in immune regulation, osteoclastogenesis, and circadian regulation. Five criteria of interactions were selected to define positive interactions: experiments (e.g., co-purification, yeast two hybrid system, and genetic interactions), database (e.g., known signaling pathways and protein complex), text-mining (e.g., unsupervised text-mining, and search for proteins frequently mentioned together), co-occurrence (e.g., gene families), and co-expression (e.g., correlated expression).

### Statistical Analysis

Unpaired two-tailed t-tests and two-way ANOVA with Tukey’s method of multiple comparisons were used as described in each figure legend. Mean ± SEM was obtained from biological replicates of the numbers indicated in each figure. Box plots show median and interquartile range (25th–75th percentile). GraphPad Prism 10 (GraphPad Software) was used in statistical analysis.

## Results

### dKO increased bone mass and decreased osteoclastogenesis in male mice

dKO and Cont mice were prepared by mating *Per1* cKO and *Per2* cKO mice. The depletion of *Per1* (23.2% ± 3.0% remains) and *Per2* (25.0% ± 4.3% remains) was verified by qRT-PCR in osteoclasts prepared from 12-week-old male mice (**Figure 1A**). Micro-CT analysis of femurs demonstrated increased bone mass in male dKO mice compared with Cont mice as shown by the increased cortical thickness (Ct.Th), trabecular bone volume (Tb.BV/TV), trabecular number (Tb.N), and connectivity density (Conn.D) (**Figure 1B and C**). dKO did not affect cortical area fraction (Ct.Ar/Tt.Ar), trabecular thickness (Tb.Th), or trabecular separation (Tb.Sp). However, the increased bone mass was not detected in female femurs (**Supplementary Figure 1**). Bone histomorphometry of the proximal tibia showed a male-specific decrease in osteoclasts per bone perimeter in dKO mice, consistent with the increased bone mass (**Figure 2A and B**, and **Supplementary Figure 2A**). The number of osteoblasts per bone perimeter was not affected by dKO in either sex (**Figure 2C and D**, and **Supplementary Figure 2B**). Therefore, we focused on male mice in the following experiments. The lack of bone phenotypes in female mice is common to *Per1* cKO.^7^

**Figure 1.**
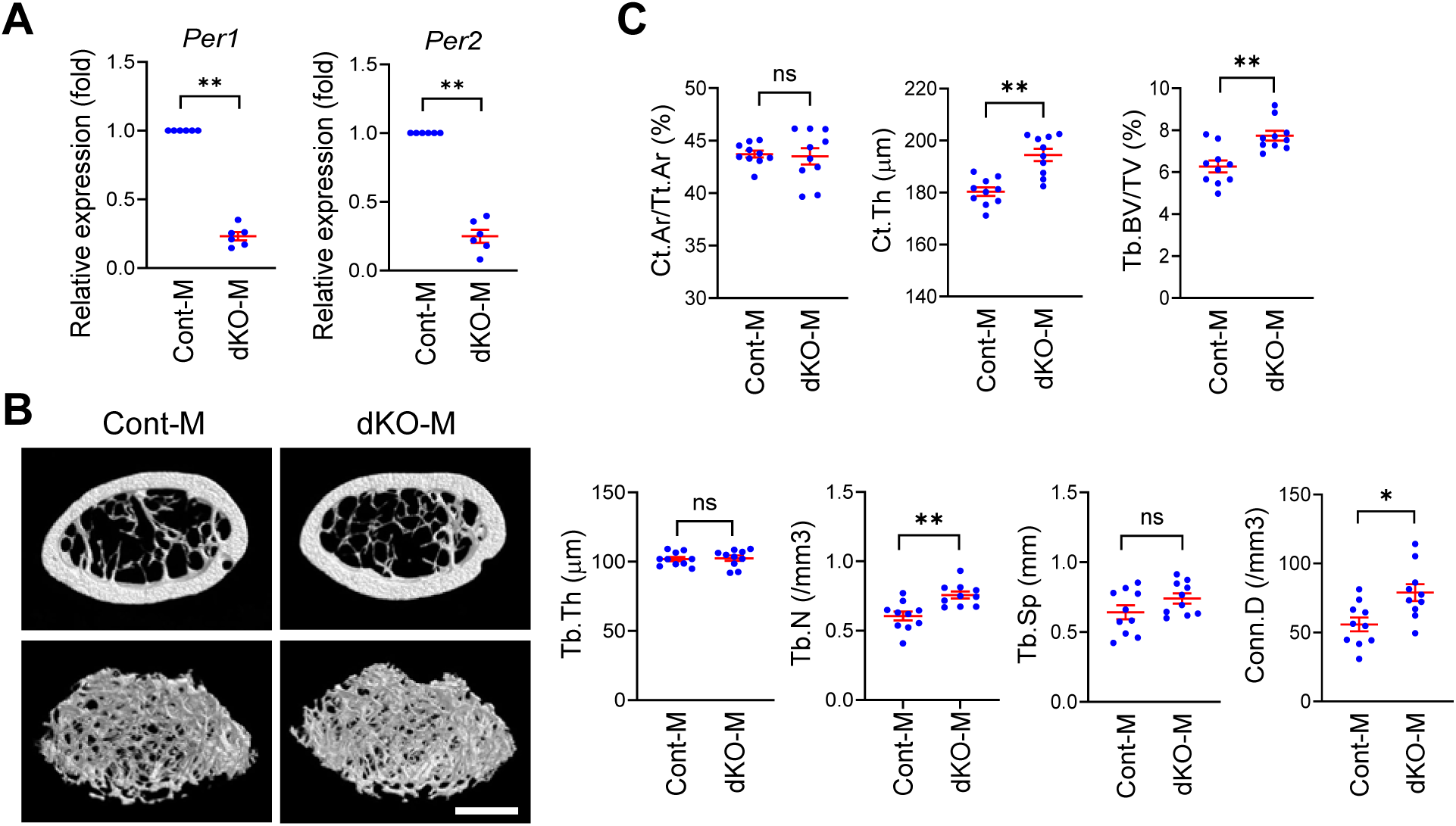
cKO increased bone mass in male femurs. **A.** Relative expression levels of *Per1* and *Per2* in dKO male osteoclasts in comparison to Cont osteoclasts on day 6 quantified with qRT-PCR. n = 6 with technical triplicates of qPCR. **B.** Reconstructed 3D micro-CT images of femoral midshafts (top) and distal femurs (bottom) prepared from 12-week-old male dKO and Cont mice. Bar, 1 mm. **C.** Quantification of cortical area fraction (Ct.Ar/Tt.Ar), cortical thickness (Ct.Th), trabecular bone volume/total volume ratio (Tb.BV/TV), trabecular thickness (Tb.Th), trabecular number (Tb.N), trabecular separation (Tb.Sp), and connectivity density (Conn.D) comparing 12-week-old male mice. n = 10. Mean ± SEM is shown. ** p < 0.01, * p < 0.05, and ns for not significant with unpaired two-tailed t-test.

**Figure 2.**
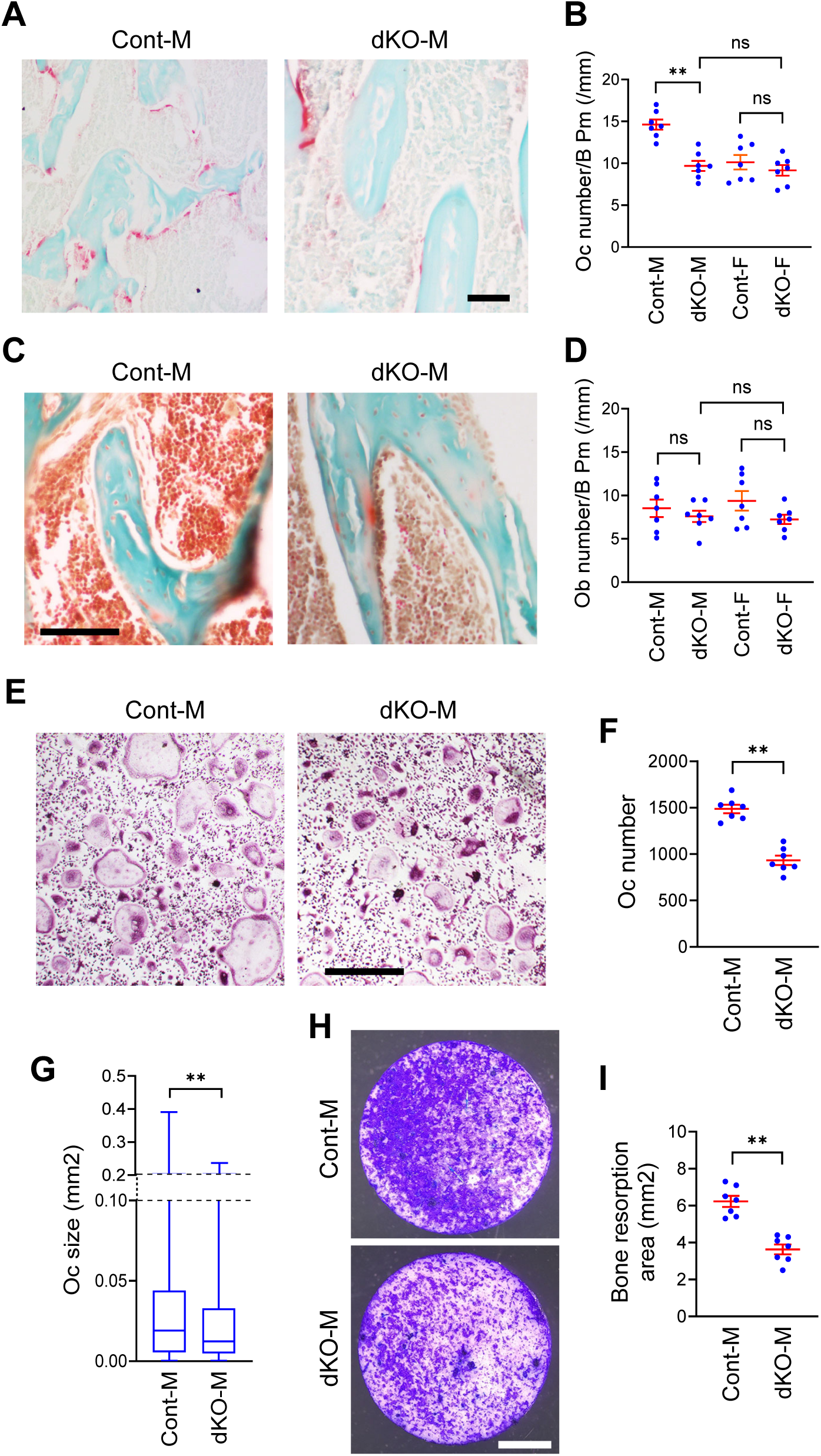
dKO increased osteoclasts in male mice. **A.** TRAP staining of the proximal tibial sections comparing dKO and Cont male mice. Bar, 100 μm. **B.** Osteoclast number per bone perimeter in the proximal tibiae of the indicated genotypes. n = 7. **C.** Masson’s trichrome staining of the proximal tibial sections. Bar, 100 μm. **D.** Osteoblast number per bone perimeter in proximal tibiae of the indicated genotypes. n = 7. **E.** TRAP staining of osteoclasts in vitro on day 6. Bar, 500 μm. **F.** The number of osteoclasts per well in a 48-well plate on day 6. n = 7. **G.** Size distribution of osteoclasts on day 6. n = 4,200 osteoclasts (600 osteoclasts/well x 7 wells of biologically independent experiments). **H.** Bone resorption assay stained with Toluidine blue. Bar, 2 mm. **I.** Areas of resorbed bones stained with Toluidine blue. n = 7. Histological sections and osteoclasts were prepared from 12-week-old male mice. Mean ± SEM is shown. ** p < 0.01, * p < 0.05, and ns for not significant with two-way ANOVA with Tukey’s method of multiple comparisons (**B** and **D**) or unpaired two-tailed t-tests (**F**, **G**, and **I**).

To understand whether the decrease in the number of osteoclasts was a cell-autonomous effect of dKO, we compared osteoclastogenesis from BMMs prepared from the femurs of dKO and Cont mice. The average number (dKO/Cont = 62.7% ± 4.6%) and size (dKO/Cont = 77.0% ± 3.2%) of osteoclasts were diminished by dKO (**Figure 2E–G**). Bone resorption was also decreased by dKO as indicated by the smaller bone resorption areas on bovine bone slices (**Figure 2H and I**). Collectively, dKO decreased osteoclastogenesis in vivo and in vitro, increasing bone mass in vivo.

### Multiple regulators of innate immunity were downregulated by dKO

RNA-seq of osteoclasts revealed that only a very small fraction of all transcripts (< 0.6%) was up-or downregulated by dKO for cutoff values of 1.5-fold difference (log2FC > 0.58 or < −0.58) and adjusted p value (padj) < 0.05 (−log10[padj] > 1.3) (**Figure 3A**). The dysregulated genes exhibited consistent patterns among biological triplicates, increasing confidence in the result (**Figure 3B**). GO analysis of the upregulated genes did not identify any enriched pathways. However, among downregulated genes, those related to innate immunity pathways were overrepresented (all top 13 pathways, including 28 genes out of 86 downregulated genes) (**Figure 3C**). They included six promoters (**Figure 3D**, blue) and six inhibitors (red) of osteoclastogenesis, in addition to 16 genes with neutral or unknown roles in osteoclastogenesis (orange). As summarized in **Table 1** (the genes marked by # are discussed later), many studies of the promoters and inhibitors did not report bone phenotypes in bone-specific cKO mice. However, in vitro phenotypes of depletion suggest that the combined effects of the counteracting genes resulted in decreased osteoclastogenesis in cKO cells. These 28 genes were not downregulated in osteoclasts with single cKO of *Per1* or *Per2* (**Supplementary Figure 3A and B**).^7^ These results indicate that all 28 genes were redundantly upregulated by *Per1* and *Per2* and that a single KO was not sufficient to substantially decrease the gene expression.

**Figure 3.**
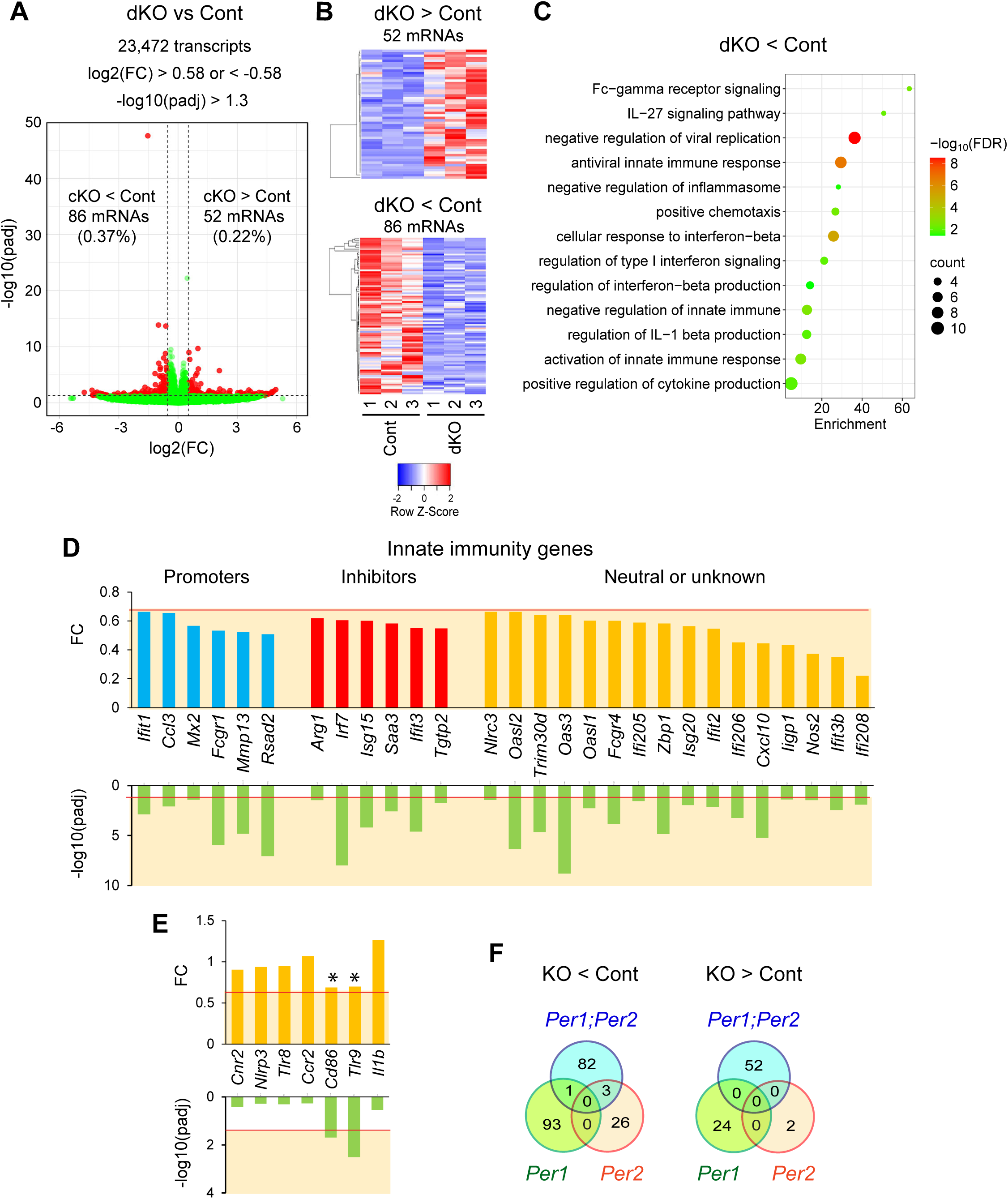
dKO downregulated innate immunity genes in male osteoclasts. **A.** A volcano plot demonstrating differentially expressed genes between dKO and Cont osteoclasts. FC and padj indicate fold change and adjusted p value, respectively. **B.** Heatmaps displaying up- or downregulated genes in dKO osteoclasts compared with Cont osteoclasts. **C.** Gene ontology analysis of downregulated genes in dKO osteoclasts compared with Cont osteoclasts. **D.** Expression levels of the innate immunity genes detected in (**C**). **E.** Expression levels of seven inflammatory genes downregulated in *Per1* KO osteoclasts. Two genes at the borderline of downregulation are indicated with asterisks. **F.** Venn diagrams displaying the numbers of up- or downregulated genes by *Per1* KO, *Per2* KO, and dKO. All data are based on n = 3 of male osteoclasts. The range of values with statistically significant difference (FC < 0.67 or –log10[padj] > 1.3) are highlighted in light orange in (**D**) and (**E**) with the cutoff values indicated by red lines. (**E**) and (**F**) used data in reference.^7^

**Table 1.**
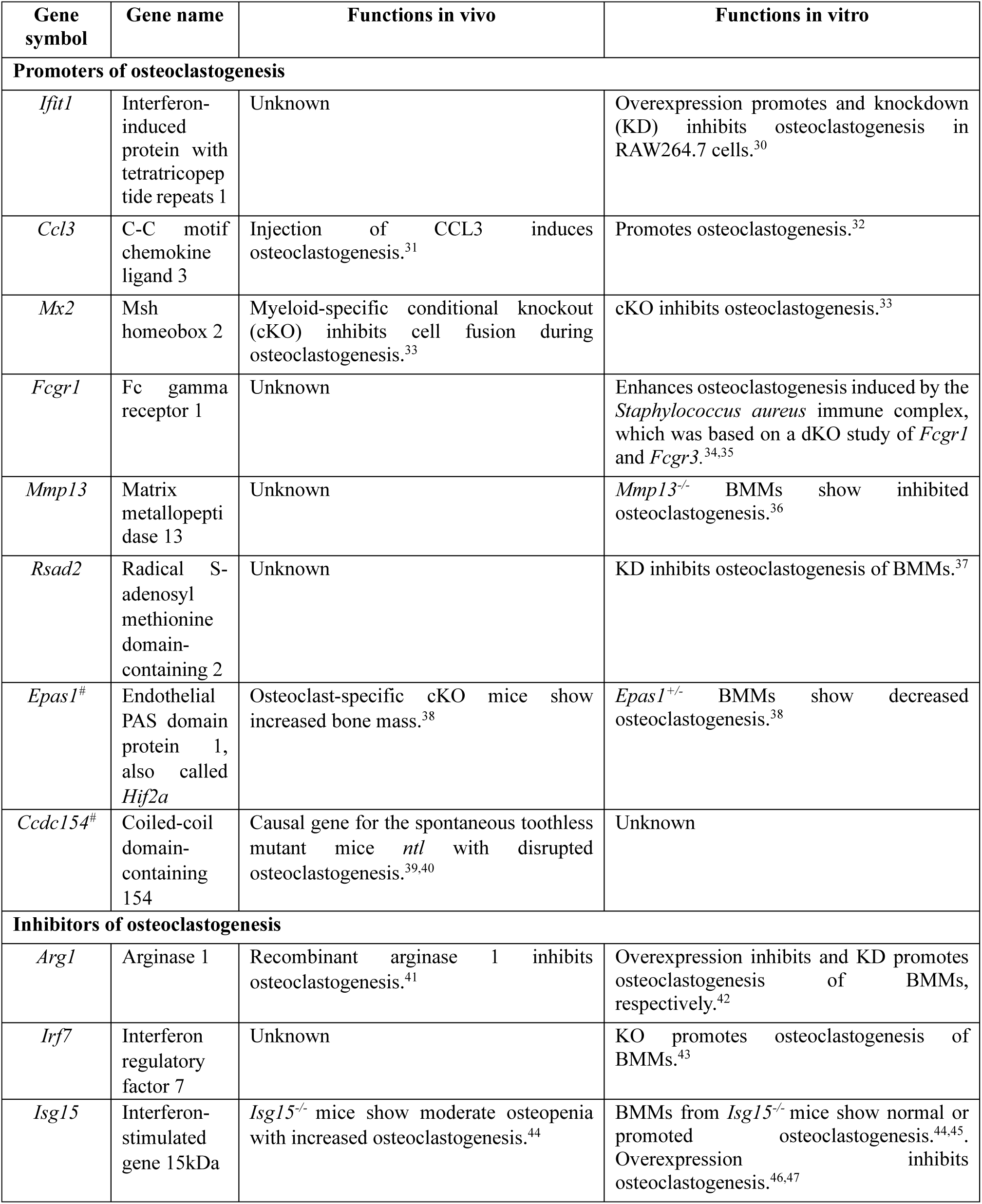
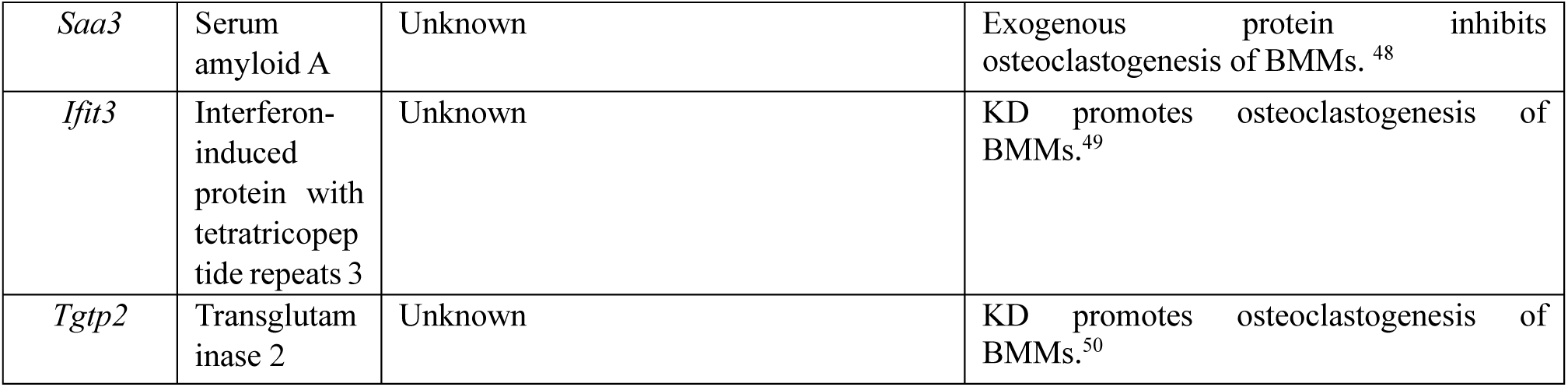
The roles of selected genes in osteoclastogenesis.

The seven inflammatory genes that were downregulated by single *Per1* KO and regulate osteoclastogenesis^7^ did not decrease in dKO cells (**Figure 3E**). This was unexpected because most genes downregulated by *Per1* KO would also have been decreased by dKO, given the lack of effects of *Per2* KO on these genes.^7^ To investigate this discrepancy, we counted the number of genes up-or downregulated by single or double KO. As shown in **Figure 3F**, there were a few overlapping genes between single and double KO cells. However, close inspection revealed that the discrepancy was more subtle than initially considered. For example, although the expression levels of *Cd86* and *Tlr8* in *Per1* KO cells did not meet our criteria of downregulation (Fold change [FC] < 0.67), their expression levels (0.69 and 0.70, respectively) were very close to the cutoff values (**Figure 3E**, asterisks). A similar situation was observed in the genes downregulated by dKO but not by *Per1* KO. Although the 12 genes marked by asterisks in **Supplementary Figure 3A** were not considered downregulated in dKO cells, their FC and padj were close to the cutoff values. Thus, the definition of cutoff values and subtle experimental variations could change the inclusion or exclusion of a given gene as up- or downregulated. These borderline genes were already downregulated to some extent by *Per1* KO and further decreased with the additional *Per2* KO, making them meet more stringent criteria of downregulation in dKO cells. In addition to this interpretation, there could be more biological reasons, such as indirect effects by dKO, to explain the discrepancy between single and double KO.

### dKO mildly downregulated numerous osteoclast-related genes

Despite clear inhibition of osteoclastogenesis in vitro, no pathways related to osteoclastogenesis emerged in the GO analysis of downregulated genes. To explore the reasons, GO analysis was repeated with less stringent cutoff values (FC < 1 and padj < 0.05). This revealed a weaker enrichment of three groups of pathways relevant to osteoclasts (osteoclast differentiation, lysosome, and actin) with some overlapping genes (**Figure 4A and B**). Osteoclasts form an actin ring to seal the resorption lacuna on the surface of bone and dissolve hydroxyapatite and bone matrix with the enzymes secreted from lysosomes, such as cathepsin k.^15^ Actin is also important to generate ruffled membrane enriched in V-ATPase, the proton pump that acidifies the resorption lacuna and promotes bone resorption. Downregulation of these genes was generally well reproduced in biological triplicates (**Figure 4C**). However, only a small number of genes met the previous cutoff values (FC < 0.67), which explains why these pathways did not appear in the previous GO analysis (**Figure 4D** and **Supplementary Figure 4A and B**. Five, three, and five genes respectively in the three groups). The 60 downregulated genes in the osteoblast differentiation group included several representative osteoclast markers, such as *Ctsk*, *Csf1r*, and *Ocstamp*, although their downregulation was mild and the biological meaning, if any, of the decrease was unclear (**Figure 4E**, purple). More importantly, *Epas1* and *Ccdc154,* whose loss disrupts osteoclastogenesis or increases bone mass, were also downregulated (**Figure 4E**, blue and **Table 1**, genes with #). *Epas1* was also at the border line of downregulation in *Per1* KO cells (**Figure 4F**, asterisk). The three genes in the actin group and five genes in the lysosome group with FC < 0.67 were not known to affect osteoclastogenesis or bone mass upon KO (**Supplementary Figure 4A and B**); they were not pursued in the current study.

**Figure 4.**
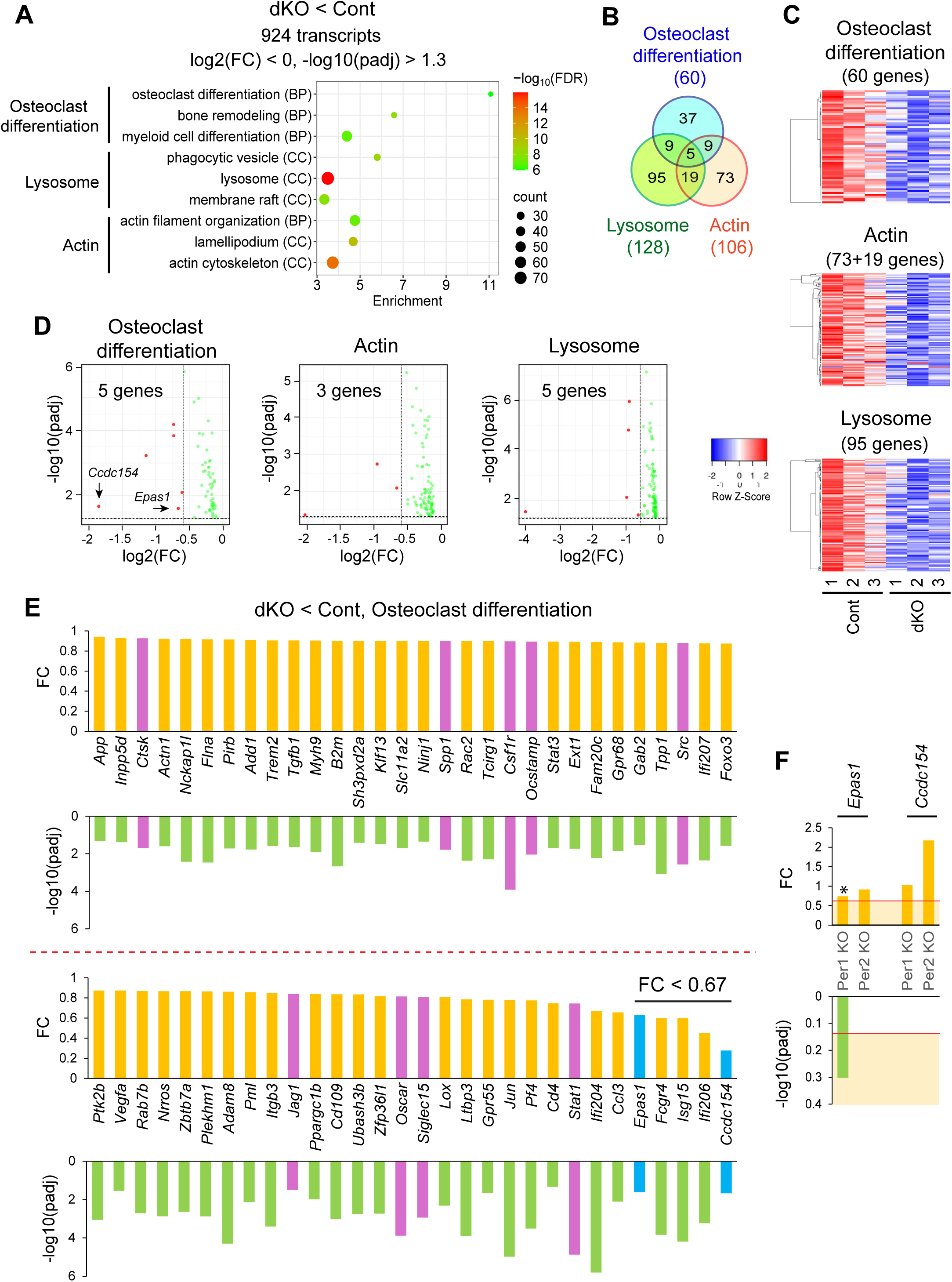
dKO mildly downregulated genes relevant to osteoclasts, actin, and lysosome. **A.** Gene ontology analysis of downregulated genes in dKO osteoclasts compared with Cont osteoclasts. Selected pathways related to osteoclast differentiation, lysosome, and actin are displayed. BP and CC indicate biological process and cellular component, respectively. **B.** A Venn diagram depicting the number of the genes in the three groups identified in (**A**) to show the number of overlapping genes. **C.** Heatmaps displaying three groups of genes downregulated in dKO osteoclasts. Actin and lysosome groups exclude the genes belonging to the osteoclast differentiation group. **D.** Volcano plots showing three groups of genes downregulated in dKO osteoclasts. The number inside each graph shows the number of genes (red dots) that met more stringent cutoff values of log2(FC) < –0.58 and –log10(padj) > 1.3. **E.** Expression levels of the genes belonging to the osteoclast differentiation group in (**A**). Representative osteoclast markers (purple) and promoters of osteoclastogenesis (blue) are highlighted. Five genes with FC < 0.67 are also indicated. **F.** Expression levels of *Epas1* and *Ccdc154* in the single KO osteoclasts of *Per1* or *Per2*. The range of values with statistically significant difference (FC < 0.67 or –log10[padj] > 1.3) are highlighted in light orange with the cutoff values indicated by red lines. The data are based on reference^7^. All data are prepared from n = 3 of male osteoclasts.

### BMAL1 binds close to some downregulated innate immunity genes

We wished to understand whether PER1 and PER2 directly regulated the genes listed in **Table 1**; however, ChIP-compatible antibodies were not currently available. As an alternative, we combined publicly available ChIP-seq data of BMAL1, our ChIP-qPCR data of BMAL1, and a reporter assay with *Per*s to investigate the possibility of direct regulation by PERs, because PERs indirectly bind chromatin via BMAL1. Most published ChIP-seq data of BMAL1 used mouse liver as material; osteoclasts or macrophage data were not available. We downloaded three pairs of input and ChIP-seq data (pairs A–C in **Figure 5A**) reported by two groups with two different BMAL1 antibodies (see the GEO database with the GSM# for more details). Four genes showed reproducible peak patterns of BMAL1 binding near or inside each gene, suggesting their direct regulation by BMAL1 (**Figure 5A**). *Saa3* and *Mx2* were selected as our model genes because of the highly reproducible BMAL1 peak patterns in three ChIP-seq data sets (**Figure 5A**, arrows).

**Figure 5.**
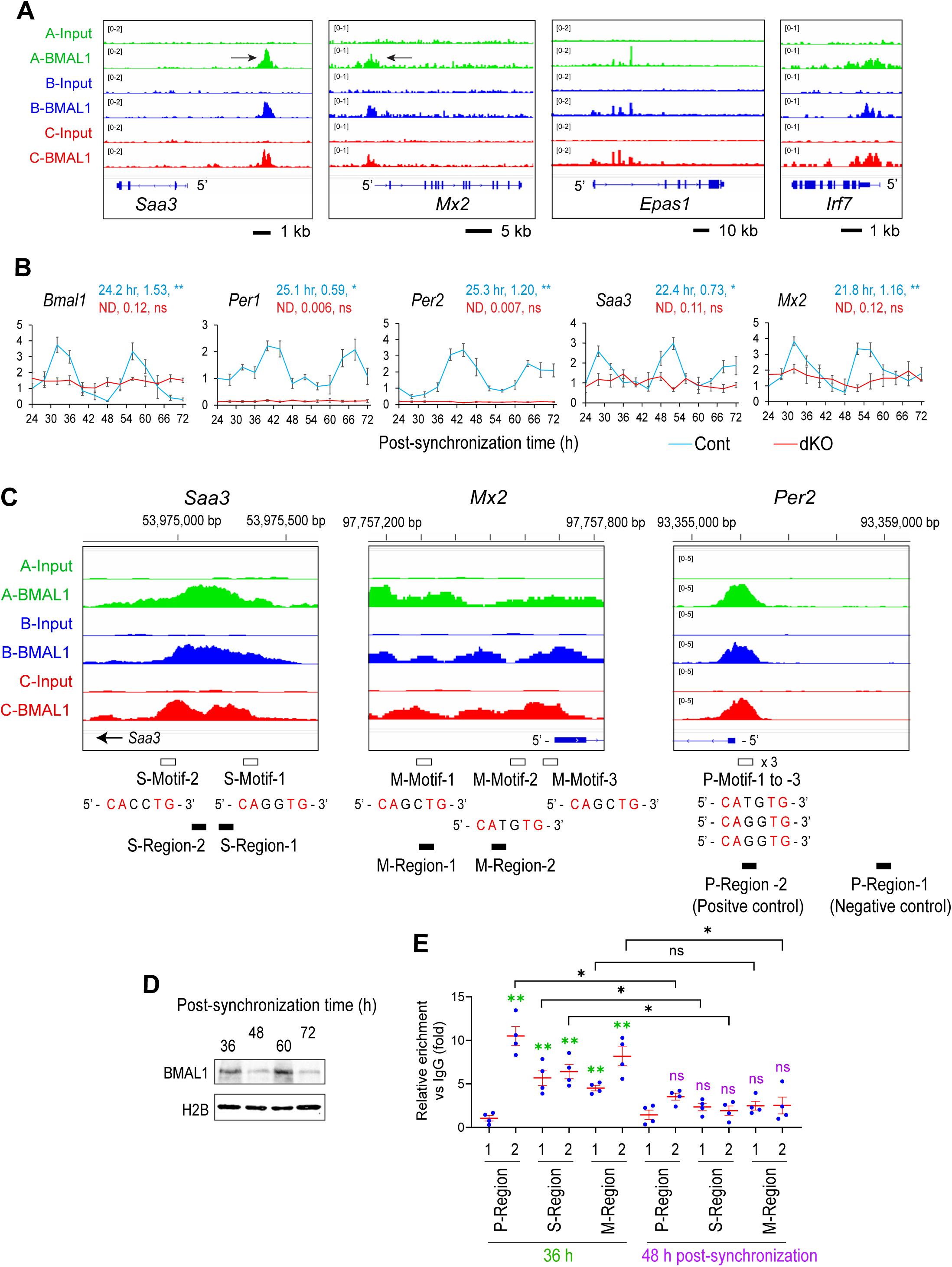
Circadian binding of BMAL1 to the *Saa3* and *Mx3* genes in osteoclasts. **A.** IGV images displaying BMAL1 peaks based on three pairs of ChIP-seq data with mouse liver downloaded from the GEO database. Arrows indicate the peaks studied in the current work. **B.** Relative expression levels of the indicated genes in synchronized osteoclasts comparing Cont and dKO osteoclasts. The value of Cont cells at 24 h was defined as 1.0 in each graph. The period (h) and amplitude of each genotype are shown above each graph. ** p < 0.01, * p < 0.05, ns for not significant, and ND for not detected with the Cosinor analysis of circadian rhythmicity above each graph. **C.** Enlarged BMAL1 peaks at *Saa3* and *Mx2* genes shown in (**A**) and at the *Per2* gene. The locations of the BMAL1-binding motifs, their nucleotide sequences, and the regions amplified by qPCR are indicated below each image. **D.** Western blotting demonstrating BMAL1 at four time points of synchronized osteoclasts. Histone H2B was used as loading control. **E.** ChI P-qPCR of BMAL1 in each region in synchronized osteoclasts. ** p < 0.01, * p < 0.05, and ns with unpaired two-tailed t-test. The statistical analysis indicated in green and purple are based on the comparison with P-Region-1 (negative control) at each time point. Male osteoclasts were used in all data. Mean ± SEM based on biological triplicates with technical triplicates are shown in (**B**) and (**E**).

First, circadian expression of the two genes was verified by synchronization of circadian rhythms of Cont osteoclasts with serum shock. *Bmal1* showed a circadian expression with a period of 24.2 h, which was lost in dKO cells as expected (**Figure 5B**). The expression patterns of *Per1* and *Per2* were anti-phasic to that of *Bmal1*, also as expected, with a period of 25.1 h and 25.3 h, respectively. The expression of *Saa3* and *Mx2* exhibited circadian patterns reaching their peak levels 6–8 h after those of *Per1* and *Per2*. This delay could be explained by the time lag required for translation and chromatin binding by PER1 and PER2. The circadian expression patterns of *Per1*, *Per2*, *Saa3*, and *Mx2* disappeared in dKO cells.

Next, we determined the DNA regions of *Saa3* and *Mx2* to be amplified by ChIP-qPCR. A close inspection of the ChIP-seq data identified two and three BMAL1-binding motifs (CANNTG) in the BMAL1 peaks near *Saa3* and *Mx2*, respectively (**Figure 5C**, S-Motifs and M-Motifs). In addition, we detected a BMAL1 peak at the 5’ end of *Per2* containing three motifs (P-Motifs), which was used as a positive control region (P-Region-2) for ChIP-qPCR because *Per2* is a well-known target gene of BMAL1 (**Figure 5C**, *Per2*). Around 3 kb upstream of *Per2* without a BMAL1 peak was selected as a negative control region (P-Region-1). ChIP primer pairs were designed to amplify two regions each within the BMAL1 peaks of *Saa3* and *Mx2* to test reproducibility (**Figure 5C**, S-Regions and M-Regions). Because the size of chromatin fragments was 200–600 bp, each PCR product could cover all motifs in each gene.

Western blotting of BMAL1 was used to decide the time of harvesting osteoclasts after synchronization. BMAL1 was abundantly expressed at 36 h and 60 h post-synchronization, whereas its level clearly decreased at 48 h and 72 h (**Figure 5D**). The abundant timings corresponded to 4 h after the peaks of mRNA, which was again consistent with the lag period due to translation.

Finally, ChIP-qPCR with BMAL1 antibody was performed comparing the cells harvested at 36 h and 48 h post-synchronization. BMAL1 was abundantly bound to the positive control region (P-Region-2) compared with the negative control region (P-Region-1) at *Per2*, validating the design of qPCR primers (**Figure 5E**). Additionally, the abundant binding of BMAL1 at the positive control region was lost at 48 h, consistent with the decreased protein level of BMAL1. BMAL1 was also bound to two regions each of *Saa3* and *Mx2* at 36 h (S-Region-1 and -2, and M-Region-1 and -2), which was lost at 48 h, reflecting the pattern at the positive control region. Collectively, these results indicate that BMAL1 was bound to the motif regions in the *Saa3* and *Mx2* genes in a circadian time-dependent manner in osteoclasts. This provided a physical basis for the study of the functional roles of BMAL1 binding in the next section.

### PER1 and PER2 activate target genes depending on the BMAL1-binding motif

To examine whether PER1 and PER2 activate the expression of *Saa3* and *Mx2*, a luciferase reporter assay was applied. A DNA fragment containing two BMAL1-binding motifs of *Saa3* was inserted into a firefly luciferase (F-Luc) expression vector as Saa3 WT (**Figure 6A**). In addition, the two motifs were mutated individually or simultaneously to create three mutants as shown in **Figure 6A**. Similarly, WT and four mutants of the three motifs of *Mx2* were created and inserted into the vector (**Figure 6B**). They were transfected into 293FT cells along with a *Renilla* luciferase (R-Luc) vector as an internal control for transfection efficiency. In addition to the reporter plasmids, *Bmal1*, *Clock*, *Per1*, *Per2*, and empty vectors (EV) were transfected in various combinations. The three mutant vectors of *Saa3* did not change the F-Luc/R-Luc ratio in the presence of EV or the combination of *Bmal1* and *Clock* (BC) compared with Saa3-WT (**Figure 6C**, EV and BC). Additional transfection of *Per1* (BCP1) or *Per2* (BCP2) increased the F-Luc/R-Luc ratio of Saa3-WT compared with BC. However, this increase was significantly inhibited by a mutation in one of the two motifs (Mut-1 and Mut-2), which was further decreased by combined mutations (Mut-12). Thus, PER1 and PER2 promoted F-Luc expression depending on both motifs. When the same experiments were repeated with *Mx2* vectors, a mutation in M-Motif-1 or M-Motif-2 diminished F-Luc expression, whereas a mutation in M-motif-3 did not affect F-Luc expression (**Figure 6D**). These results indicate that PER1 and PER2 use only the first two motifs in *Mx2*.

This interpretation was verified by an independent approach by transfecting *Per1* and *Per2* into the macrophage-derived cell line RAW264.7 along with *Bmal1* and *Clock*. BC did not upregulate *Saa3* or *Mx2* compared with the EV; however, BCP1 and BCP2 could upregulate the genes (**Figure 6E**). *Per1* or *Per2* alone could not upregulate *Saa3* or *Mx2*, which aligned with the known dependence of PER1 and PER2 on BMAL1 and CLOCK for gene regulation (**Figure 6E**). The endogenous *Bmal1* and *Clock* appeared insufficient to support the roles of *Per*s.

**Figure 6.**
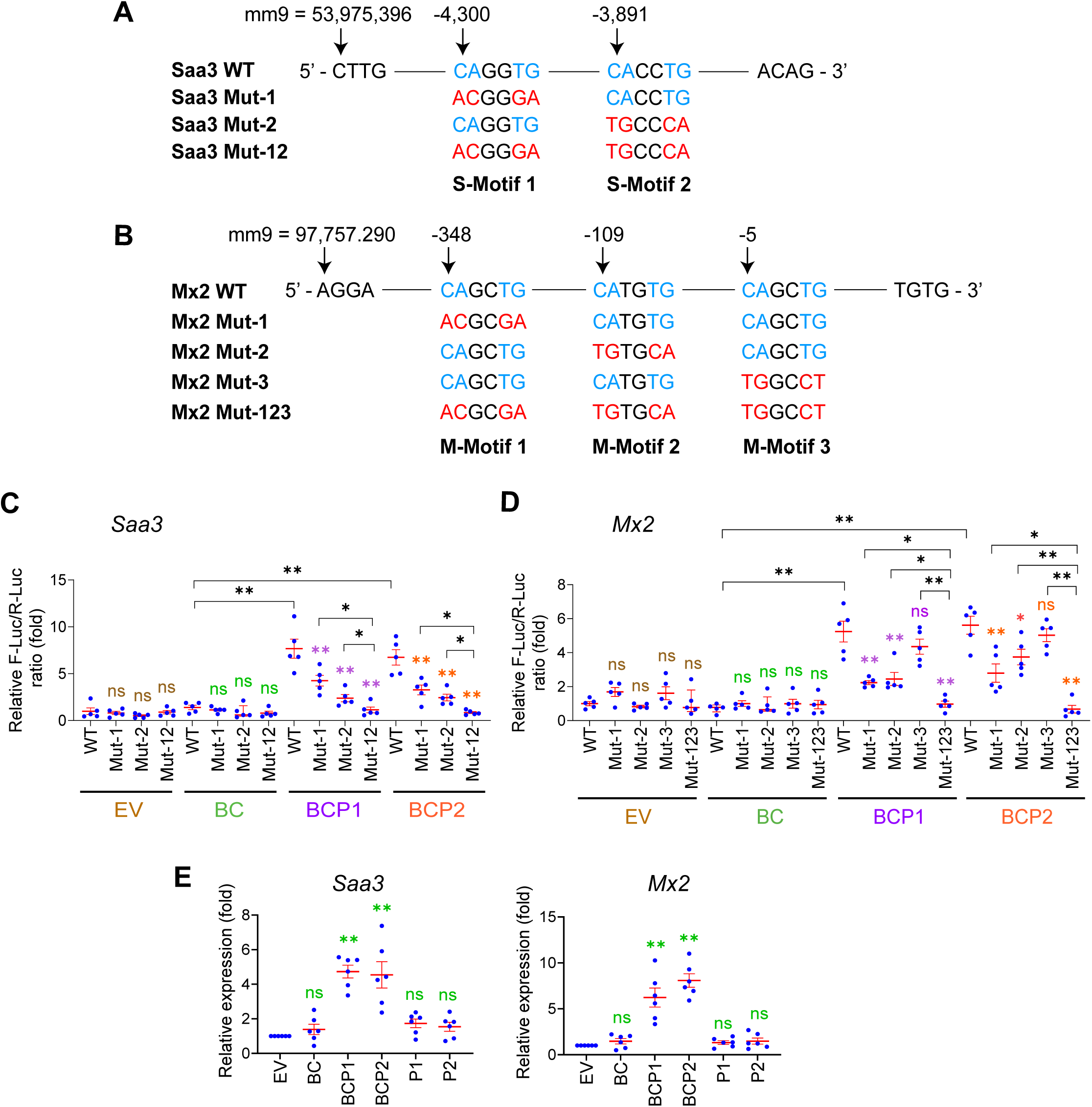
Activation of the *Saa3* and *Mx2* genes by *Per1* and *Per2*. **A** and **B.** Sequences of the BMAL1-binding motifs in the wild type (WT) and mutants (Mut) sequences *Saa3* (**A**) and *Mx2* (**B**). Conserved (light blue) and mutated (red) nucleotides are indicated in color. mm9 numbers indicate the nucleotide numbers based on NCBI37/mm9. The minus numbers above the initial C in each motif represent the nucleotide numbers counted from the transcription start site. **C** and **D.** Relative F-Luc/R-Luc ratios after transfection of the *Saa3* (**C**) and *Mx2* (**D**) reporter plasmids into 293FT cells. Results with EV (empty vector), BC (*Bmal1* and *Clock*), BCP1 (BC with *Per1*), BCP2 (BC with *Per2*) are shown with the value with WT and EV defined as 1.0 in each graph. **E.** Relative expression levels of *Saa3* and *Mx2* upon transfection of the indicated gene combinations into RAW264.7 cells. The value with EV was defined as 1.0 in each graph. Mean ± SEM based on biological triplicates with technical triplicates are shown in (**C**)–(**E**). ** p < 0.01, * p < 0.05, and ns for not significant with unpaired two-tailed t-test in comparison to WT in each gene combination as color-coded in (**C**) and (**D**) or in comparison to EV in (**E**). Additional comparisons are shown with black lines in (**C**) and (**D**).

### Per1/Per2 integrate interactomes of circadian regulators, innate immunity genes, and osteoclast genes

These studies functionally linked three groups of genes—circadian rhythms, osteoclastogenesis, and innate immunity. To understand already known interactomes between their genes, the STRING program was applied to the 86 genes downregulated in dKO osteoclasts (**Figure 3A**), 60 mildly downregulated osteoclast genes (**Figure 4E**, five overlapped with the 86 genes), and 11 circadian regulators. The program defined interactions from experimentally verified specific cases to simple co-expression in databases to frequent co-occurrence in literature. Three major clusters emerged from this analysis. First, there was a network of genes relevant to innate immunity (**Figure 7**, surrounded by the purple dashed line). This tightly integrated promoters (blue circle) and inhibitors (red circle) of osteoclastogenesis together with genes without known effects on osteoclastogenesis (green circle). The second cluster (surrounded by the orange dashed line) also included innate immunity genes but most were regulators of osteoclast differentiation or functions (white circles). These two clusters intersected at *Cxcl10* and *Stat3* as shared members with unknown roles in osteoclasts. In addition, there were 46 osteoclast genes (white circles) outside or loosely connected to the two clusters. The third cluster—circadian regulators (surrounded by the pink dashed line)—was largely segregated from the other two clusters and surrounding isolated osteoclast genes. There was no direct link between Per1/Per2 (underlined in red) and the genes outside the circadian cluster. *Bmal1* (underlined in red) interacted with *Epas1* (connected with a red line), a promoter of osteoclastogenesis; however, this interaction was based on a variant of yeast two-hybrid system without functional investigation.^16^ Our study linked all these genes under a large umbrella of circadian regulation, with *Per1* and *Per2* as key mediators.

**Figure 7.**
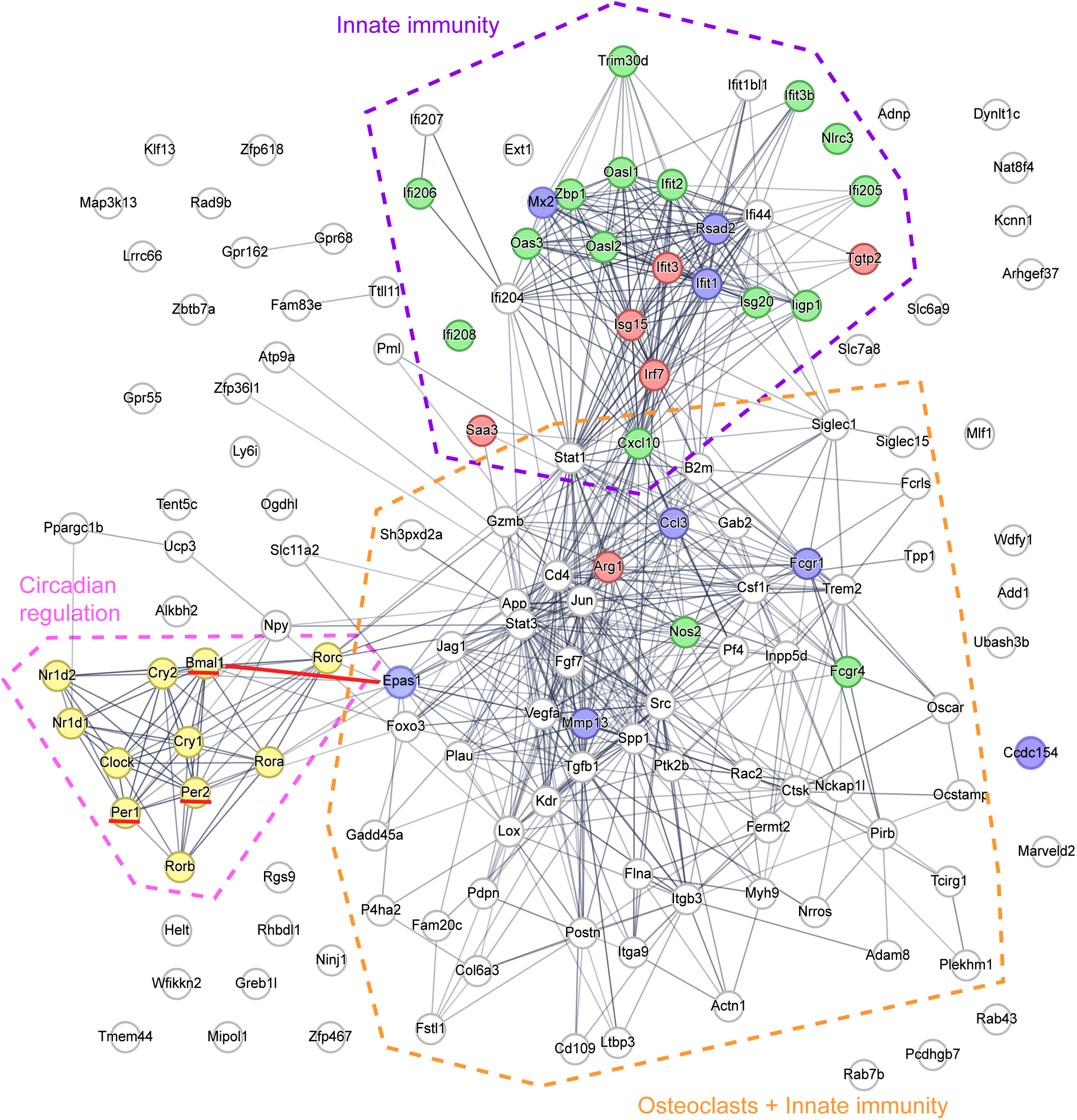
Network analysis of the genes involved in innate immunity, osteoclasts, and circadian regulation. Color code of each circle is as follows. Innate immunity genes that promote osteoclastogenesis (blue), inhibit osteoclastogenesis (red), and are neutral or with unknown roles in osteoclastogenesis (green). Circadian regulators (yellow). Osteoclast genes (white).

## Discussion

This study showed that *Per1*;*Per2* dKO inhibited osteoclastogenesis and increased bone mass in mice. However, in our previous work, single KO of *Per1* had opposite effects—promoted osteoclastogenesis and decreased bone mass.^7^ Furthermore, single KO of *Per2* does not affect bone mass or osteoclastogenesis. Thus, KO of the two paralogs, which share 73% similarity at the amino acid level, demonstrated a spectrum of bone phenotypes depending on the context. Each KO dysregulated distinct sets of genes with some overlap, consistent with the variability in phenotypes. Nonetheless, preferential downregulation of genes related to various immune reactions emerged as a common trend in dKO and *Per1* KO osteoclasts. There was no single overwhelmingly dominant pathway involving the genes. The downregulated genes included promoters and inhibitors of osteoclastogenesis. Therefore, the bone phenotypes caused by *Per1* and/or *Per2* KO represent combined effects of multiple competing activities in osteoclastogenesis. Some appear directly controlled by PERs through chromatin binding. Thus, PERs can be considered multifaceted controllers of osteoclastogenesis, rather than pure promoters or inhibitors. The broad effects of *Per* KO are hardly surprising, given the widespread gene regulation by circadian rhythms: 26% of the transcriptome in the mouse calvaria are under circadian influence.^17^

The circadian fluctuation of immunological genes in osteoclasts is a part of a wider perspective on the gene regulation in the monocyte/macrophage lineages, where expression of immunological genes intrinsically fluctuates in circadian manners without stimuli.^18^ Consequently, the intensity of the cell’s immune reactions to external stimuli depends on the circadian timing of the stimulation. For example, macrophages exhibit circadian expression of a variety of inflammatory signaling genes, such as toll-like receptors and TNF-α pathway genes.^19,20^ The magnitude of the response of macrophages to lipopolysaccharide (LPS) is markedly differs depending on the time of day of the administration of LPS, providing functional relevance to the gene expression patterns.^19,21^ Similar circadian patterns in immunological genes have been reported in other cells within the monocyte/macrophage lineages, such as microglia in the brain, Kupffer cells in the liver, and alveolar macrophages in the lung.^22–24^ Moreover, circadian regulation of immunity is also evident beyond the monocyte/macrophage lineages, such as B cells, T cells, dendritic cells, and neutrophils.^18^ Circadian activity of the immune system is thought to be an underlying cause of the daily fluctuation in the joint stiffness and pain in rheumatoid arthritis patients, in addition to hormonal rhythms.^2^ It is possible that circadian changes in immunological gene expression in osteoclasts also contribute to a circadian fluctuation of the symptoms. This is impossible to test in patients because of a lack of easy access to bone marrow cells. However, our findings open a future research direction focusing on the dysregulation of innate immunity genes as a possible link between inflammatory bone diseases and circadian rhythms.

Compared with macrophages and the tissue-specific counterparts mentioned above, evidence for the control of immunological genes by specific circadian regulators is scarce in osteoclasts. Studies of osteoclast-specific *Bmal1* KO did not report any changes in immunological genes.^25,26^ Little is known about the phenotypes and gene expression in the osteoclast-specific cKO mice of other circadian regulators.^2^ In this context, our finding placed all genes marked by red, blue, green, and white circles in **Figure 7** under the direct or indirect influence of PER1 and PER2, expanding the scope of the impact of specific circadian regulators.

Epidemiological studies have shown a correlation between shift work and an increased risk of osteoporosis and fracture, likely due to multiple factors, such as sleep deprivation, hormonal disruption, irregular meals, and insufficient physical exercise.^27,28^ Although shift works up- or downregulate circadian regulators in white blood cells, the causal relationship with the increased risk of osteoporosis remains unknown. It would be inappropriate to directly infer a causal implication of our findings for shift workers’ osteoporosis; however, serum inflammatory cytokines are elevated in shift workers without apparent underlying diseases.^20,29^ In addition, mice maintained under shifted light-dark cycles demonstrate elevated serum proinflammatory cytokines with and without stimulation by LPS.^20^ Thus, despite the different triggers (*Per* KO vs shifted light cycle), immunological disruption is a common outcome, supporting the use of KO mouse models to the study of shift work-induced osteoporosis.

## Supporting information

Graphical abstract

## Acknowledgments

We are grateful to the Comparative Pathology Shared Resource and the Minnesota Dental Research Center for Biomaterials and Biomechanics.

## Author contributions

Nobuko Katoku-Kikyo (investigation, writing - review and editing), Elizabeth K. Vu (investigation, writing - review and editing), Samuel Mitchell (investigation, writing - review and editing), Ismael Y. Karkache (investigation, writing - review and editing), Elizabeth W. Bradley (conceptualization, funding acquisition, investigation, writing - review and editing), and Nobuaki Kikyo (conceptualization, funding acquisition, investigation, writing - original draft, writing - review and editing).

## Data Availability

RNA-seq data have been deposited to Gene Expression Omnibus (GEO) under the accession number of GSE293417.

## Funding Statement

E.W.B. was supported by the NIH (R21AR084530). N.K was supported by the NIH (R01GM137603) and Regenerative Medicine Minnesota (RMM 072523 DS 002). The content is solely the responsibility of the authors and does not necessarily represent the official views of the NIH.

## Conflict of interest

The authors declare no conflict of interest.

## Notes

### Competing Interest Statement

The authors have declared no competing interest.

