## Supplementary Materials for "Circadian regulators PER1 and PER2 regulate osteoclastogenesis by balancing competing activities of innate immunity genes"

[*] Corresponding authors

**List of Supplementary Material**

Supplementary Figure 1–4

Supplementary Figure Legends

Supplementary Table 1 and 2

Supplementary Fig 1

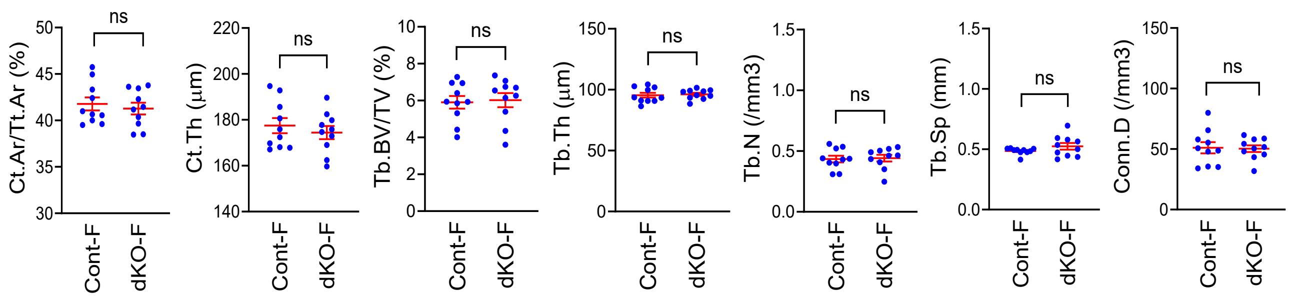

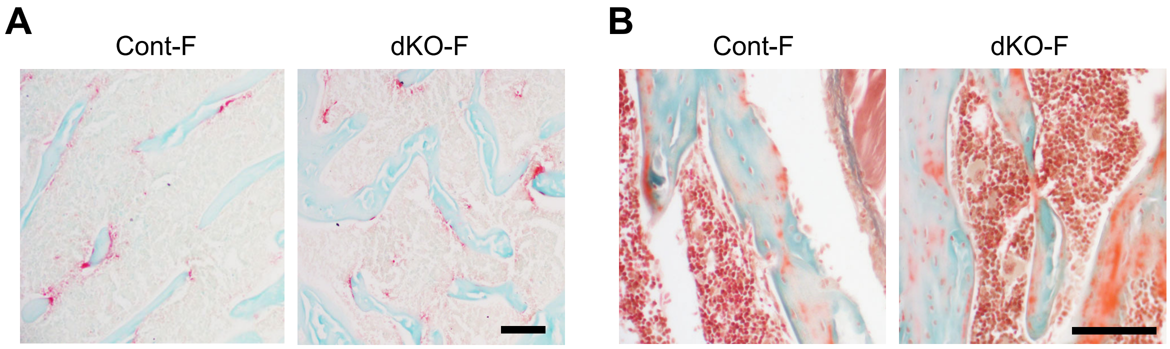
A
Cont-F
dKO-F
B
Cont-F
dKO-F

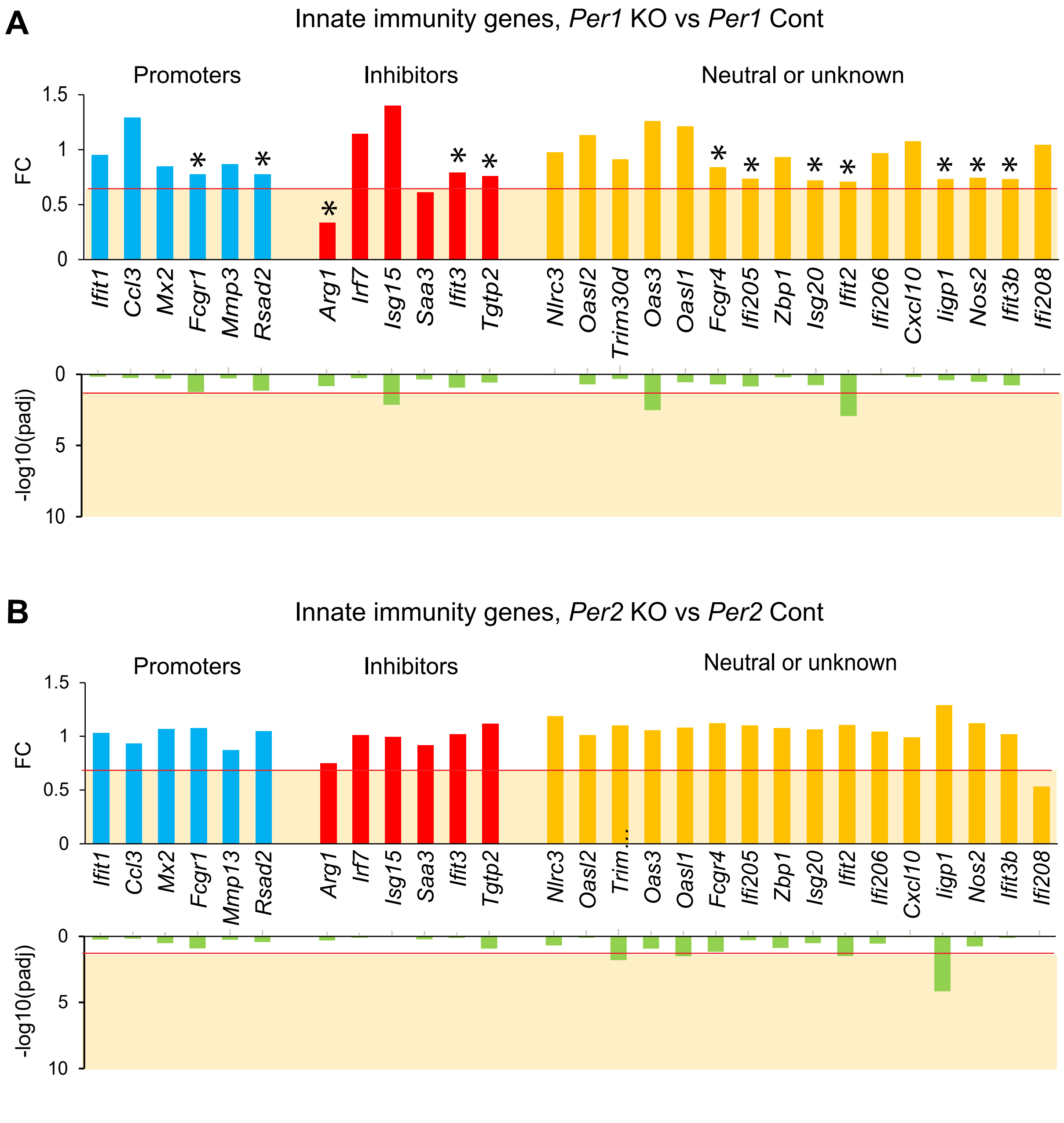
A
Innate immunity genes, Per1 KO vs Per1 Cont
Promoters
Inhibitors
Neutral or unknown
FC
-log10(padj)
Ifit1
Ccl3
Mx2
Fcgr1
Mmp3
Rsad2
Arg1
Irf7
Isg15
Saa3
Ifit3
Tgtp2
Nlrc3
Oasl2
Trim30d
Oas3
Oasl1
Fcgr4
Ifi205
Zbp1
Isg20
Ifit2
Ifi206
Cxcl10
Iigp1
Nos2
Ifit3b
Ifi208
B
Innate immunity genes, Per2 KO vs Per2 Cont
Mmp13
Trim…

Supplementary Fig 4

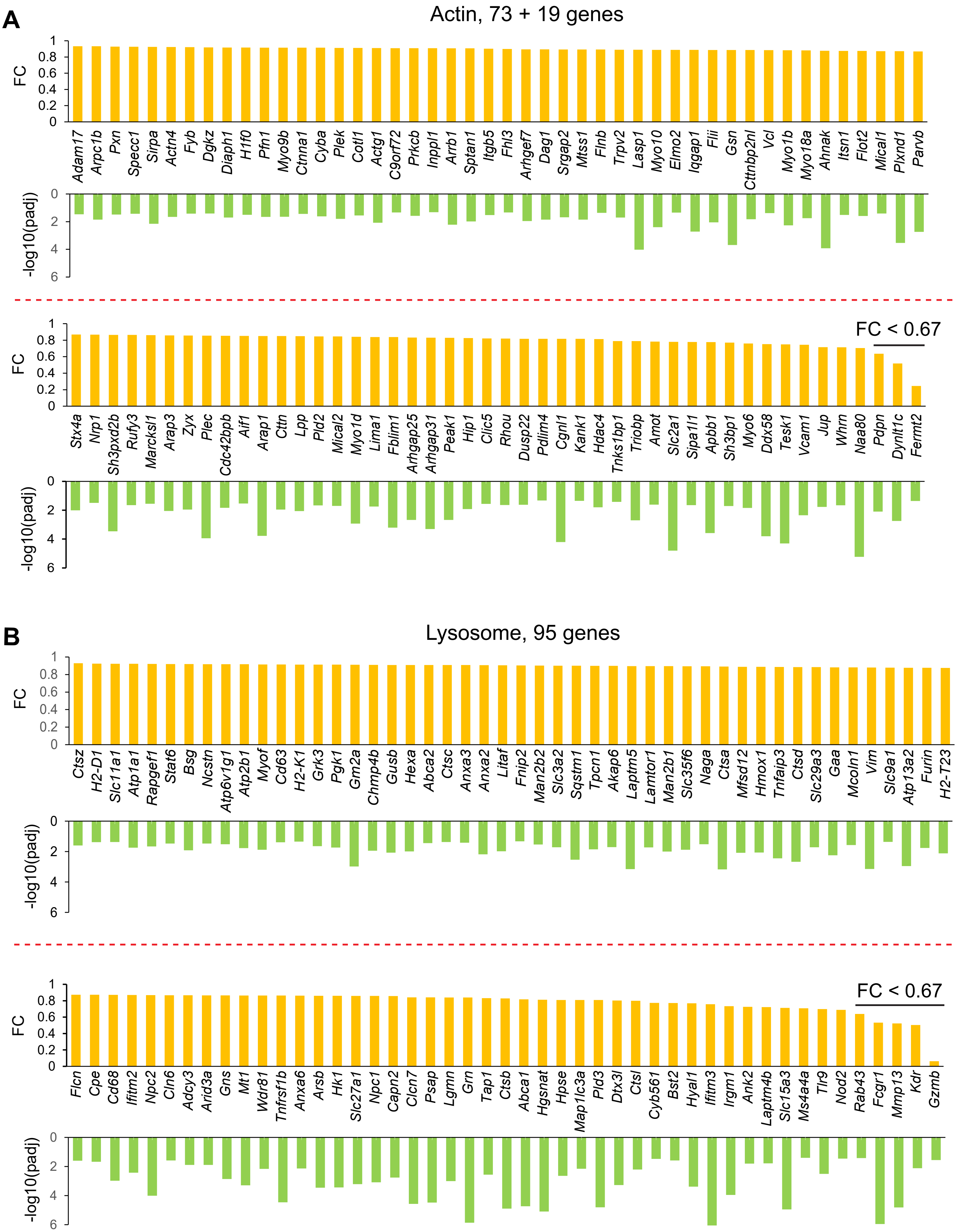

### Supplementary figure legends

**Supplementary Figure 1. Bone mass of femurs was not affected in female dKO mice.**

Quantification of cortical area fraction (Ct.Ar/Tt.Ar), cortical thickness (Ct.Th), trabecular bone volume/total volume ratio (Tb.BV/TV), trabecular thickness (Tb.Th), trabecular number (Tb.N), trabecular separation (Tb.Sp), and connectivity density (Conn.D) comparing 12-week-old female mice of dKO and Cont. Mean ± SEM from n = 10 mice. ns for not significant with unpaired two-tailed t-test.

**Supplementary Figure 2. Osteoclastogenesis and osteoblastogenesis were not affected in dKO female mice.**

**A.** TRAP staining of the proximal tibial sections comparing dKO and Cont mice.

**B.** Masson's trichrome staining of the proximal tibial sections.

Histological sections were prepared from 12-week-old female mice. Bar, 100 μm.

**Supplementary Figure 3. Expression levels of selected innate immunity genes in *Per1* KO and *Per2* KO osteoclasts.**

**A** and **B.** Expression levels of the innate immunity genes downregulated in dKO osteoclasts. The comparison between *Per1* KO and Cont (**A**) and *Per2* KO and Cont (**B**) are shown. All data are based on n = 3 of male cells. The range of the values with statistically significant difference (FC [fold change] < 0.67 or –log10[padj] > 1.3) are highlighted in light orange with the cutoff values indicated by red lines. The data are based on reference[7].

**Supplementary Figure 4. Downregulated genes related to actin and lysosome in dKO**

**osteoclasts.**

**A** and **B.** Expression levels of 73 genes unique to the actin group and 19 genes overlapping between actin and lysosome groups (**A**) and 95 genes unique to the lysosome group (**B**) in dKO osteoclasts. The genes with FC < 0.67 are highlighted. All data are based on n = 3 male cells.

**Supplementary Table 1. Sequences of qRT-PCR primers**

| Gene | Forward | Reverse |
|---|---|---|
| *Gapdh* | TGCACCACCAACTGCTTAG | GATGCAGGGATGATGTTC |
| *Bmal1* | CAACCCATACACAGAAGCAAAC | CATCTGCTGCCCTGAGAATTA |
| *Per1* | CCTGGAGGAATTGGAGCATATC | CCTGCCTGCTCCGAAATATAG |
| *Per2* | CAAAGCTGACGCACACAAAG | TTAGCCTTCACCTGCTTCAC |
| *Saa3* | GCCTTCCATTGCCATCATTC | CACATGTCTCTAGACCCTTGAC |
| *Mx2* | CTACTGCCAGGACCAGATTTAC | GTGCCATGCTTTGTCTTCTTC |

**Supplementary Table 2. Sequences of ChIP-qPCR primers**

| Gene | Forward | Reverse |
|---|---|---|
| S-Region-1 | GAACGTGCTGTGCTGTATTTAG | AATTGACCCTCCCAGAGATTAC |
| S-Region-2 | CTAGGCTCAGTACCATCCAAAC | CAAGGAGCTTACCTGCTGAA |
| M-Region-1 | AAGGCCAGCACTTCTCAT | AAGGGTCAGAGACAAACCATAC |
| M-Region-2 | CCCTCTAAACTGGTGTCCATAC | CTCAGAAAGGAAACTCACCCTAA |
| P-Region-1 (negative control) | GGAGACTGAGGTGACTTTCTTG | GGGCTGATCTCACTATCCTCTA |
| P-Region-2 (positive control) | GGAGTTCCATGTGCGTCTTAT | TGCCACCTCATTTGCATACT |
