## Supplementary material for "Circadian regulators PER1 and PER2 regulate osteoclastogenesis by balancing competing activities of innate immunity genes": Graphical abstract

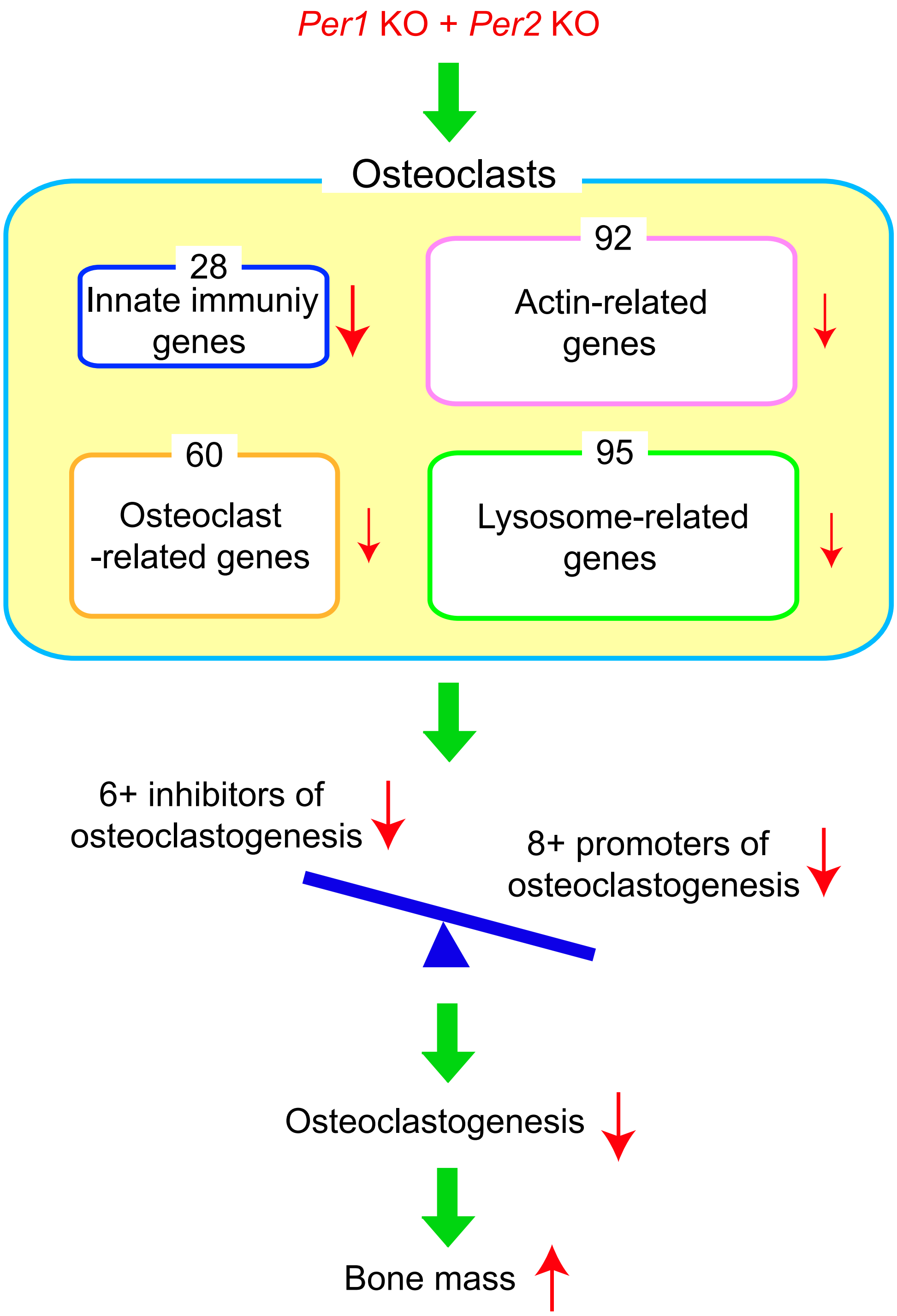

**Graphical abstract**

Double knockout of the circadian regulator paralogs *Per1* and *Per2* downregulates > 200 genes in mouse osteoclasts, most prominently innate immunity gene and less severely the genes related to osteoclasts, actin, and lysosome. They include multiple established promoters and inhibitors of osteoclastogenesis. The combined effects result in decreased osteoclastogenesis and increased bone mass. This work integrates circadian regulators, innate immunity, and osteoclastogenesis.
